# Deep learning for inferring gene relationships from single-cell expression data

**DOI:** 10.1101/365007

**Authors:** Ye Yuan, Ziv Bar-Joseph

## Abstract

Several methods were developed to mine gene-gene relationships from expression data. Examples include correlation and mutual information methods for co-expression analysis, clustering and undirected graphical models for functional assignments and directed graphical models for pathway reconstruction. Using a novel encoding for gene expression data, followed by deep neural networks analysis, we present a framework that can successfully address all these diverse tasks. We show that our method, CNNC, improves upon prior methods in tasks ranging from predicting transcription factor targets to identifying disease related genes to causality inference. CNNC’s encoding provides insights about some of the decisions it makes and their biological basis. CNNC is flexible and can easily be extended to integrate additional types of genomics data leading to further improvements in its performance.

**Supporting website with software and data:** https://github.com/xiaoyeye/CNNC.

## Introduction

Several computational methods have been developed to infer relationships between genes based on gene expression data. These range from methods for inferring co-expression relationships between pairs of genes^1^ to methods for inferring a biological or disease process for a gene based on other genes (either using clustering or guilt by association^2^) to causality inferences^3, 4^ and pathway reconstruction method^5^. To date, each of these tasks was handled by a different computational framework. For example, gene co-expression analysis is usually performed using Pearson correlation or mutual information^6^. Functional assignment of genes is often performed using clustering^7^ or undirected graphical models including Markov random fields^8^, while pathway reconstruction is often based on directed probabilistic graphical models^4^. These methods also serve as an initial step in some of the most widely used tools for the analysis of genomics data including network inference and reconstruction approaches^3, 9, 10^, methods for classification based on genes expression^11^ and many more.

While successful and widely used, these methods also suffer from serious drawbacks. First, each relies on (different) manually determined assumptions about the distribution of the observed values. Another major issue is overfitting. Most of these methods are unsupervised. Given the large number of genes that are profiled, and the often relatively small (at least in comparison) number of samples, several genes that are determined to be co-expressed or co-functional may only reflect chance or noise in the data^12^. Finally, most of the widely used methods are symmetric which means that each pair has only one relationship value. While this is advantageous for some applications (for example, clustering) it may be problematic for methods that aim at inferring causality (for example, network reconstruction tasks).

To address these issues we developed a new method, CNNC which provides a supervised way (that can be tailored to the condition / question of interest) to perform gene relationship inference. CNNC utilizes a novel representation of the input data specifically suitable for deep learning. It represents each pair of genes as an (image) histogram and uses convolutional neural networks (CNNs) to infer relationships between different expression levels encoded in the image. The network is trained with positive and negative examples for the specific domain of interest (for example, known targets of a TF, known pathways for a specific biological process, known disease genes etc.) and the output can be either binary or multinomial.

We applied CNNC using a large cohort of single cell expression data and tested it on several inference tasks. We show that CNNC outperforms prior methods for inferring interactions (including TF-gene and protein-protein interactions), causality inference, functional assignments (including biological processes and disease), and as a component in algorithms for the reconstruction of known pathways.

## Results

We developed CNNC a general computational framework for supervised gene relationship inference (**Fig. 1**). CNNC is based on a CNN which is used to analyze summarized co-occurrence histograms from pairs of genes in scRNA-Seq data. Given a relatively small labeled set of positive pairs (with either negative or random pairs serving as negative) CNNC learns to discriminate between interacting, causal pairs, negative pairs or any other gene relationship types that can be defined.

**Figure 1:**
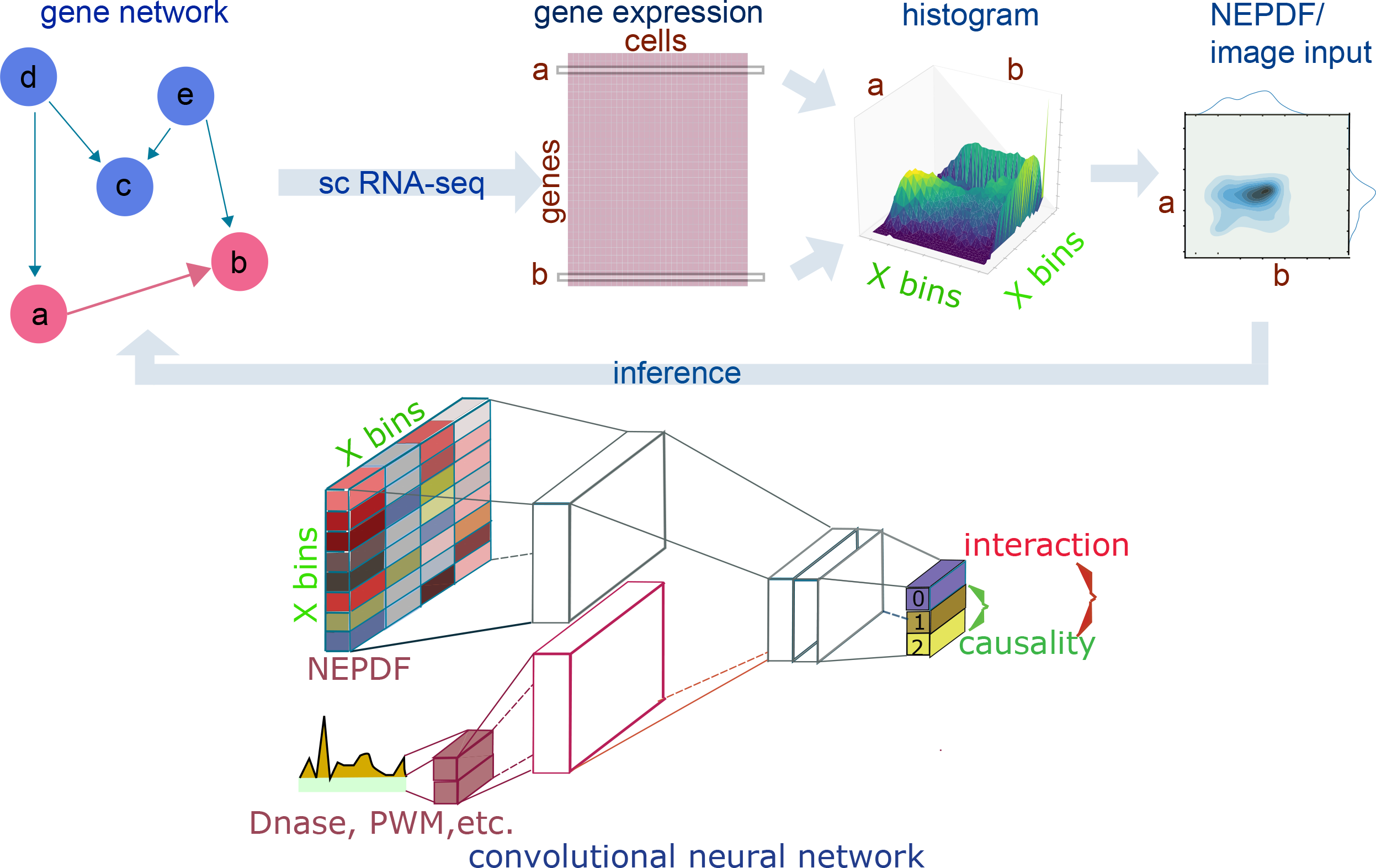
CNNC input and architecture. CNNC aims to infer gene-gene relationships using single cell expression data. For each gene pair, sc RNA-Seq expression levels are transformed into 32×32 normalized empirical probability function (NEPDF) matrices. The NEPDF serves as an input to a convolutional neural network (CNN). The intermediate layer of the CNN can be further concatenated with input vectors representing Dnase-seq and PWM data. The output layer can either have a single, three or more values, depending on the application. For example, for causality inference the output layer contains three probability nodes where *p0* represents the probability that genes *a* and *b* are not interacting, *p1* encodes the case that gene *a* regulates gene *b*, and *p2* is the probability that gene *b* regulates gene *a*.

### Learning a CNNC model

CNNC can be trained with any expression dataset, though as with other neural network applications the more data the better its performance. Given expression data we first generate a normalized empirical probability distribution function (NEPDF) for each gene pair (genes *a* and *b*) (**Fig. 1**). For this we calculate normalized 2-dimension (2D) histogram of fixed size (32×32), where columns represent gene *a* expression levels and rows represent gene *b* such that entries in the matrix represent the (normalized) co-occurrences of these values. If different data types are combined (for example, Bulk and SC) they can be either used separately or concatenated to form a combined NEPDF with dimension of 32×64. Next, the distribution matrix is used as input to a CNN which is trained using a N-dimension (ND) output label vector, where N depends on specific task. For example, for co-expression or interaction prediction N is set to 1 (interacting or not) while for causality inference it is set to 3 where label 0 indicates that genes *a* and *b* are not interacting and label 1 (2) indicates that gene *a* (*b*) regulates gene *b* (*a*). In general, our CNN model consists of one 32×32 input layer, ten intermediate layers including six convolutional layers, three maxpooling layers, one flatten layer, and a final ND ‘softmax’ layer or one scalar ‘Sigmoid’ layer (**Methods** and **Supplementary Fig. 1**).

For the analysis presented in this paper we used processed scRNA-Seq data of 43,261 cells that was collected from over 500 different studies representing a wide range of cell types, conditions etc^13^. All raw data was uniformly processed and assigned to a pre-determined set of more than 20,000 mouse genes (**Methods**).

In addition to gene expression data, CNNC can integrate other data types including Dnase-seq^14^, PWM^15^, etc. For this, we concatenated the additional information as a vector to the intermediate output of the gene expression data and continued with the standard CNN architecture. See **Methods** and **Supplementary Fig. 1** for complete details and **Supplementary Table 1** for information on training and run time.

### Using CNNC to predict TF-gene interactions

We first tested the CNNC framework on the task of predicting pairwise interactions from gene expression data^16^. Chromatin immunoprecipitation (ChIP)-seq has been widely used as a gold standard for studying cell-specific protein-DNA interactions^17^. We thus evaluated CNNC’s performance using cell-type specific scRNA-seq datasets (three for mouse embryonic stem cells (mESC), and one each for bone marrow and dendritic cells, **Methods**) and ChIP-seq data from GTRD^18^.

We extracted data from GTRD for 38 TFs for which ChIP-seq experiments were performed in mESC, 13 TFs studied in bone marrow cells and 16 TFs for dendritic cells. To determine targets for each TF using the ChIP-seq data we followed prior work^19, 20^ and defined a promotor region as 10KB upstream to 1KB downstream from the transcription start site (TSS) for each gene. If a TF X has at least one detected peak signal in or overlapping the promotor region of gene Y, we say that TF X regulates gene Y. For this prediction task we compared CNNC with several popular methods for gene-gene co-expression analysis: Pearson correlation (PC) and mutual information (MI) that are the two most popular co-expression analysis methods, Genie3^9^ which was the best performer in the DREAM4 In Silico networks construction challenge and Count statistics^21^ which relies on local information based on gene expression ranks in large heterogeneous samples. Since prior methods used for comparison are symmetric, we focused here on the two labels setting (interacting or not). We performed leave-one-TF-out cross validation analysis. For each dataset, we trained CNNC with all other TFs and used the left-out TF for testing (**Methods**).

**Fig. 2** presents the results of these comparisons. As can be seen, CNNC outperforms all prior methods for all cell types. We observe significant improvement over all prior methods (**Fig. 2k**). The AUROC achieved by CNNC is around 40% higher than PC and close to 25% higher for MI on some datasets (See **Supplementary Fig. 2** for details). Importantly, as can be seen in **Fig. 2a-2e**, the difference is even more pronounced for the top ranked predictions. For CNNC we see almost no false negatives for the top 15% ranked pairs. Such top predictions are often the most important since the ability to validate predicted interaction is usually limited to the top few predictions.

**Figure 2.**
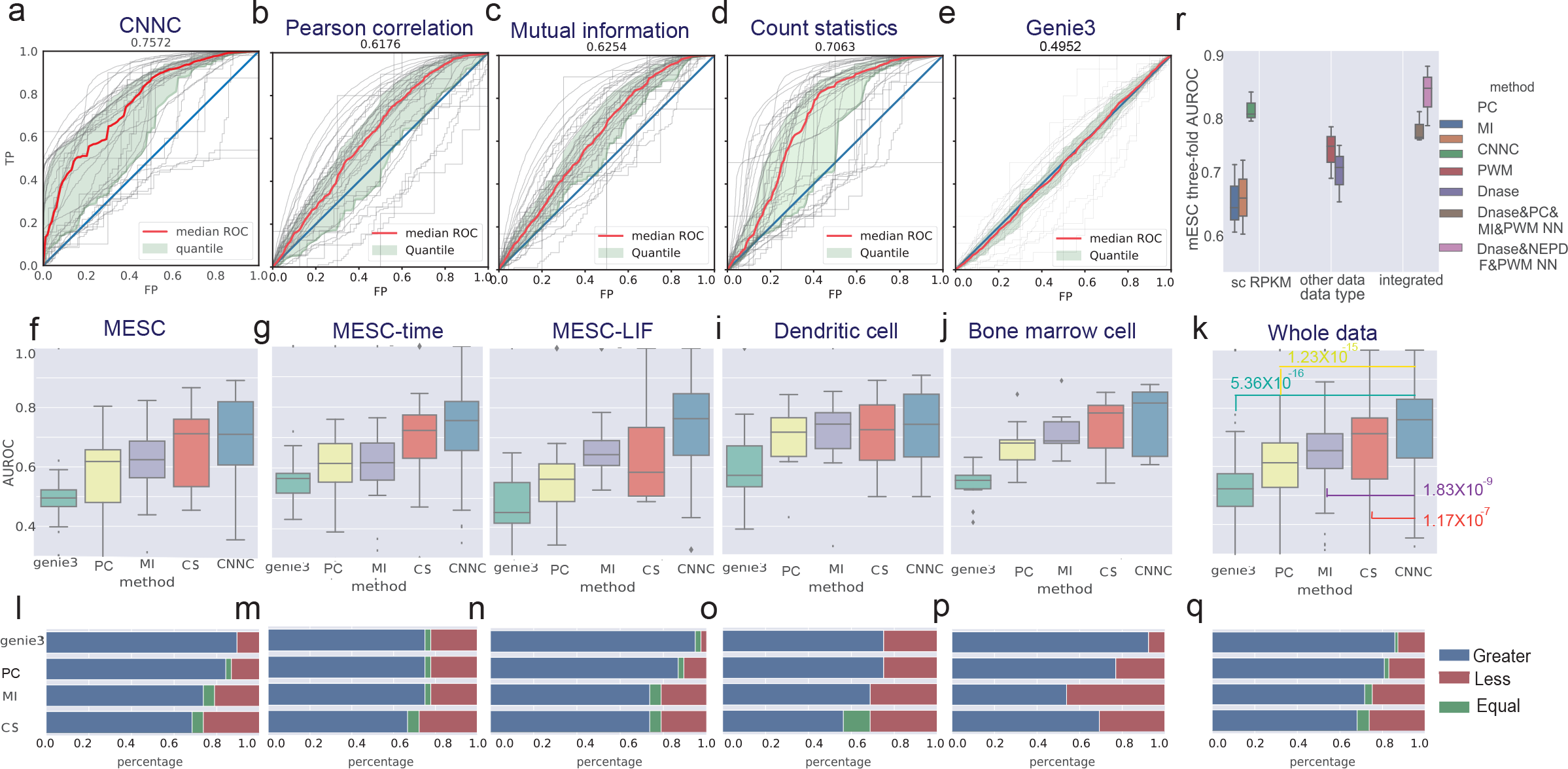
GTRD TF-target prediction. (**a-e**) ROCs of CNNC, Pearson correlation (PC), Mutual information (MI), Count statistics (CS) and Genie3 trained on scRNA-Seq mESC expression data. We performed cross validation using 38 TFs that were profiled in the same cell type. Light gray lines represent the performance for each TF. Red line represents the median ROC, and light green region represents the 25~75 quantile. (**f-k**) Result summary for the five methods using additional scRNA-Seq expression input sets and TFs, (**k**) the summary for the whole five datasets, where P-value is based on wilcoxon test comparing CNNC to all prior methods. (**l-q**) Percentage of TFs in which CNNC improves upon all other methods, (**q**) the result for the whole five datasets. (**r**) Comparison of TF-target predictions with additional data using mESC expression and TFs. Columns 1-3 show median AUROC of PC, MI, and CNNC using scRNA-Seq data respectively. 4^th^ and 5^th^ column show performance when only using PWM or Dnase. The last two columns show performance of the integration of expression, sequence (PWM) and DNase data.

### Data Integration further improves TF target gene prediction

The above analysis was only based on using expression values. However, as noted above, gene relationship inference is often used as a component in more extensive procedures that often integrate different types of genomics data. To test how the use of the NN-based method can aid such procedures we extended CNNC so that it can utilize sequence and DNase hypersensitivity information. For sequence, we used PWMs from Jaspar^22^. Dnase-seq data for mESC was obtained from the mouse ENCODE project^23^. We used a simple strategy for processing the PWM and DNase data which resulted in an additional 2D vector as input for each pair which we embedded to create a 512D vector (**Methods**). We next concatenated this vector with the NEPDF’s 512D vector in the flatten layer to form a 1024D vector as shown in **Fig. 1** and **Supplementary Fig. 1**.

Results, presented in **Fig. 2p**, show that these additional data sources indeed improve the ability to predict TF-gene interactions. As before, a combined framework utilizing CNNC outperforms a method that used both MI and PC.

### CNNC can predict pathway regulator-target gene pairs

While TFs usually directly impact the expression of their targets, several methods have also utilized RNA-Seq data to infer pathways that combine protein-protein and protein-DNA interactions^24^. To test whether CNNC can serve as a component in pathway inference methods we selected two representative pathway databases, KEGG^25^ and Reactome^26^ as gold standard and used these, together with a large scRNA-Seq dataset^13^ to train and test our framework. Since we are interested in causal relationships we only used directed edges with activation or inhibition edge types and filtered out cyclic gene pairs where genes regulate each other mutually (to allow for a unique label for each pair). As for the negative data, here we limited the negative set to a random set of pairs where both genes appear in pathways in the database but do not interact. Given the large number of genes we performed a three-fold cross validation where we kept the set of genes for which we predicted interactions completely separated (so a gene in the test set does not have any interaction in the training set, **Methods**). Results are presented in **Fig. 3**. As can be seen, CNNC performs very well on the KEGG pathways reaching an AUROC of 0.97 compared to less than 0.87 for the methods we compared which here also included Bayesian Directed Networks (BDN)^4^ which learn a global directed interaction graph, (Fig. 3b) (See **Supplementary Fig. 3** for the different folds). CNNC also performs well on Reactome pathways (see **Supplementary Fig. 3 and 4**). We also used the KEGG data to test the specific architecture CNNC utilizes and observed that the architecture used improves upon two alternative deep NN architectures, fully connected NN and CNN without pooling layers (**Supplementary Fig. 5)**.

**Figure 3.**
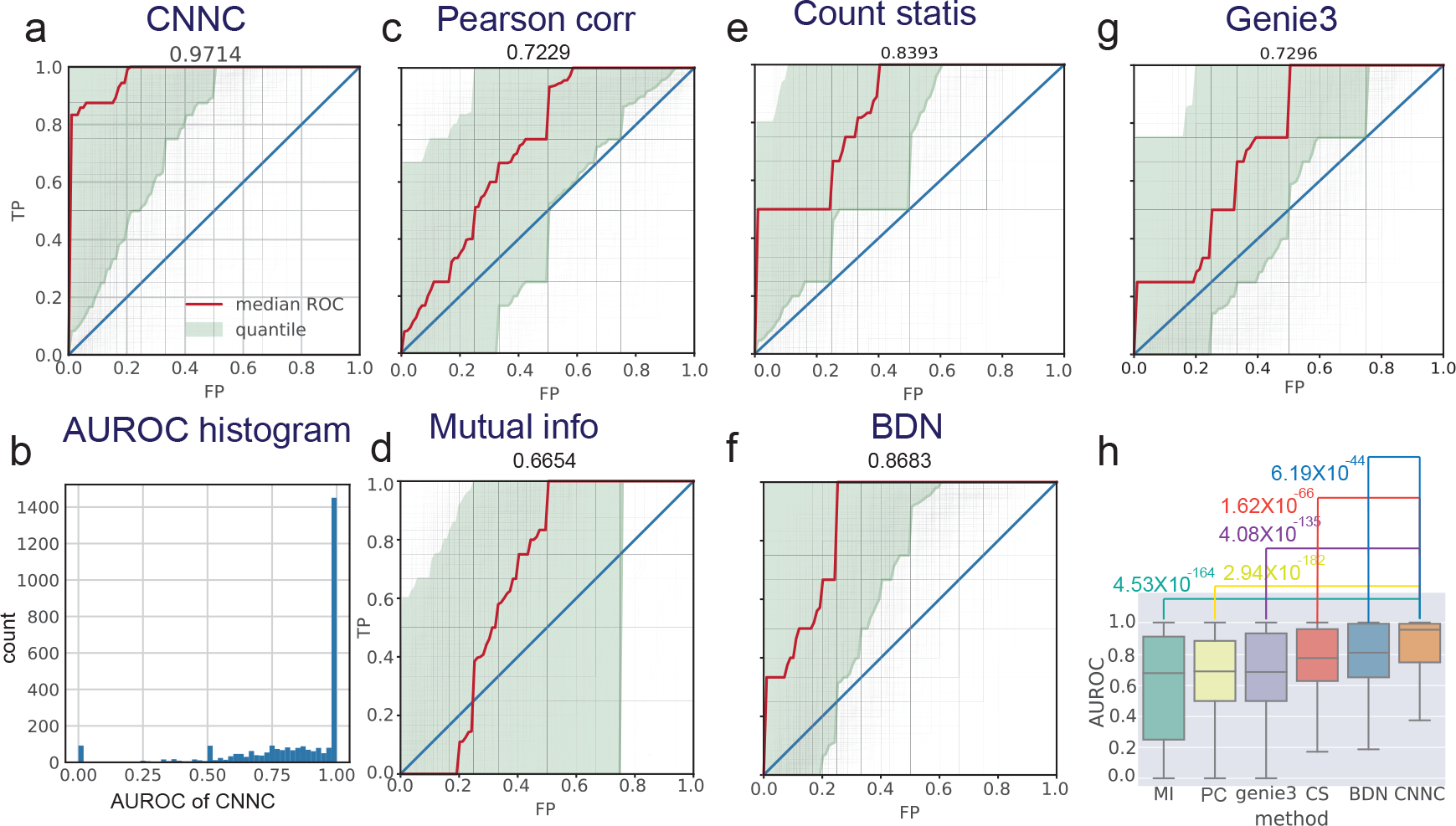
Predicting undirected pathway edges. (**a**) Overall ROCs for CNNC performance on KEGG pathway gene interaction prediction using a large compendium of scRNA-Seq data and bulk data. (**b**) The Area Under the Receiver Operating Characteristic curve (AUROC) histogram for (**a**). (**c-g**) Overall ROCs for Pearson correlation, mutual information, count statistics, Bayesian directed network (BDN) and Genie3 when tested on the KEGG pathway gene interaction prediction task. (**h**) Comparison of the six methods on the gene interaction prediction task. P-values are based on wilcoxon test. Boxplot was shown with median, first, third quartile, maximum and minimum.

### Using CNNC for causality prediction

So far we focused on general interaction predictions. However, as discussed above CNNC can also be used to infer directionality by changing the output of the NN. We next used CNNC to infer causal edges for all the datasets above (TF, KEGG and Reactome). For the pathway databases we only analyzed directed edges and so had the ground truth for that data as well. As can be seen in **Fig. 4**, when using the TF GTRD dataset, CNNC achieves a median AUROC of 0.9342 (**Fig. 4a)** on this leave-one-TF-out classification task. See **Supplementary Fig. 6** for other datasets results on this task. For KEGG, CNNC is very successful achieving a median AUROC of 0.9949 (**Fig. 4c**) (See **Supplementary Fig. 6** for the different folds). For Reactome (**Fig. 4e**) we see that the most confident predictions are correct, but beyond the top prediction performance levels off (See **Supplementary Fig. 6** for the different folds). We compared the performance of CNNC to another method developed for learning causal relationships from gene expression data, BDN^4^ which learns a global directed interaction graph. Results presented in **Supplementary Fig. 7** and **8** show that CNNC greatly outperforms BDN on this causality prediction task. We have also tested an application of CNNC that in addition to causality can infer the impact of the interaction (activation or repression) and determined that it performs well on this multi-label classification problem as well (See **Supplementary Fig. 9**).

**Figure 4.**
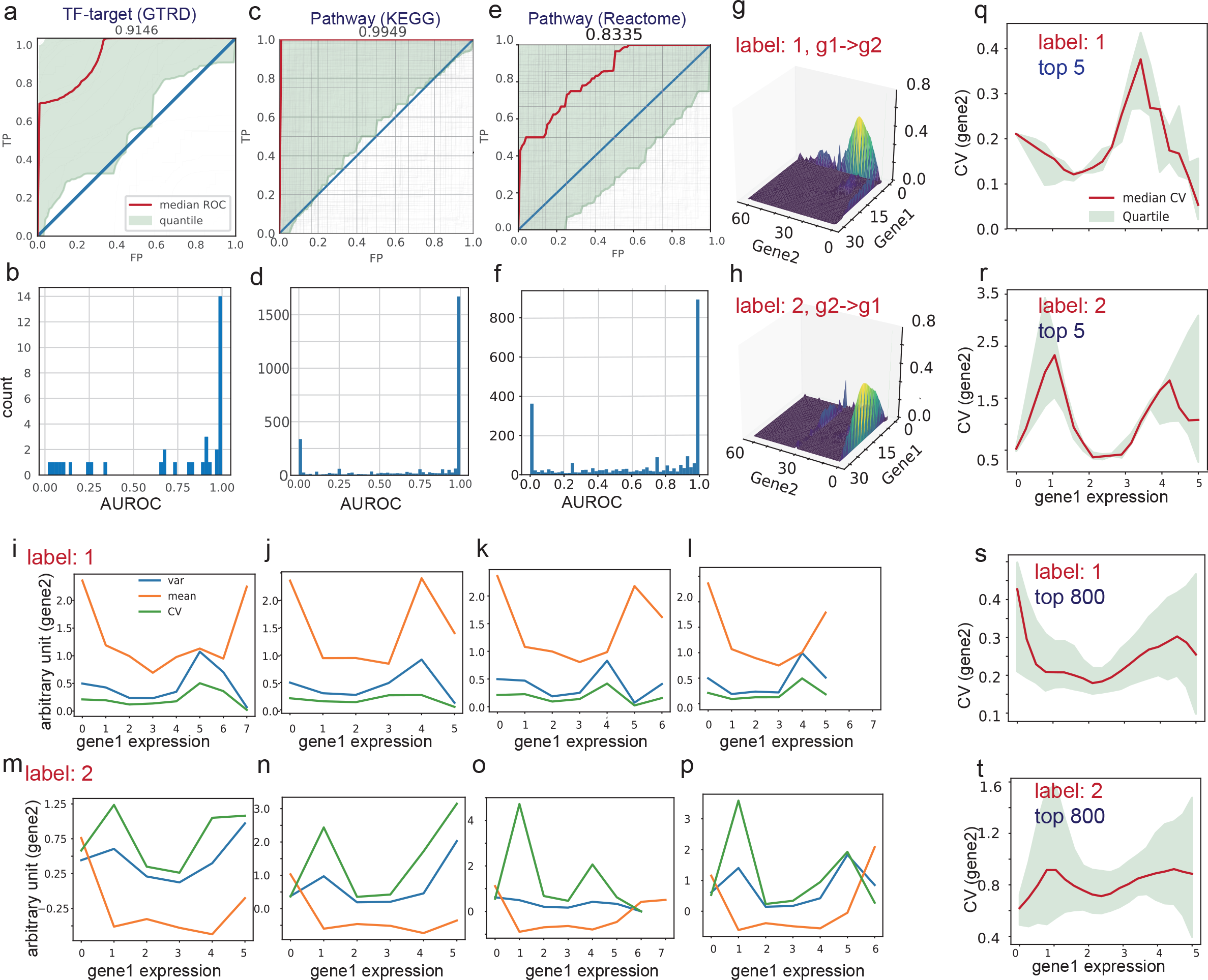
Directed (causal) edge prediction. (**a**) Overall ROCs for performance of CNNC on GTRD directed prediction task using the mESC-time dataset and mESC TFs. (**b**) The AUROC histogram for (**a**). (**c**) Overall ROCs for performance of CNNC on KEGG pathway directed edge prediction using a large compendium of scRNA-Seq and bulk data. (**d**) The AUROC histogram for (**c**). (**e**) Overall ROCs for performance of CNNC on Reactome pathway directed edge prediction. (**f**) The AUROC histogram for (**e**). (**g**) A typical NEPDF sample from a KEGG interaction that is correctly predicted as label 1. (**h**) A typical NEPDF sample that is correctly predicted as label 2. (**i-l**) Variance (var), mean and coefficient of variance (CV) of gene 2 as the expression of gene 1 increases for top correctly predicted pairs with label 1. (**m-p**) Same for top predictions for label 2. (**q-r**) Average and variance of CV for the top prediction groups correctly predicted as label 1 (**q, s**) and label 2 (**r,t**).

To try to understand the basis for the decisions reached by CNNC we plotted two of the NEPDF inputs (**Figs. 4g and 4h**) which were correctly predicted as two different labels (1 for **4g** and 2 for **4h**). As can be seen, in both inputs the two genes display partial correlations and there are places where both are up or down concurrently. However, the main difference between the histograms in **4g** and **4h** are cases where one gene is up and the other is not. In **4g** gene 2 is up while gene 1 is not indicating that the causal relationship is likely g1 -> g2. The opposite holds for **4h** and so the method infers that g2 -> g1 for that input. While relationships between expression values, including the ones mentioned above, can be manually prescribed for an algorithm, we also noted that the encoding used for CNNC allows it to look at more complicated relationships between genes. In **Figure 4i-p** we plot the mean, variance and coefficient of variance (CV) for gene2 as a function of the expression of gene 1 for both prediction directions (1->2, top and 2->1, bottom). As can be seen, the variance and CV trends are consistent within category and diverging between categories, indicating that CNNC can make use of second order or even higher order distribution properties. Similar phenomena has been anecdotally observed in specific cases, for example for miRNA regulation^27^ but the ability of CNNC to learn such relationships on its own strongly suggests that it can generalize much better than prior methods for inferring such causal interactions.

### Using CNNC for functional assignments

We next explored the use of CNNC for assigning function or disease relevance for genes. We started by using it to identify cell cycle genes. For this, we obtained 682 cell cycle genes from GSEA^28^. Since cell cycle expression has been studied for over two decades we expect most cell cycle genes to be known and so we can treat these genes as ground truth, unlike for several other processes and diseases. We next trained CNNC using all expression data on 2/3 of these genes holding the other 1/3 as a test set. In this setting the network is trained to predict 1 for a pair of genes that are both cell cycle genes and 0 for all other pairs (**Methods and supplementary Note**). When testing on the held out set CNNC achieved a high AUROC of 0.82. Importantly, the top 20% predicted genes were all true positives (**Fig. 5a** and **5b**).

**Figure 5.**
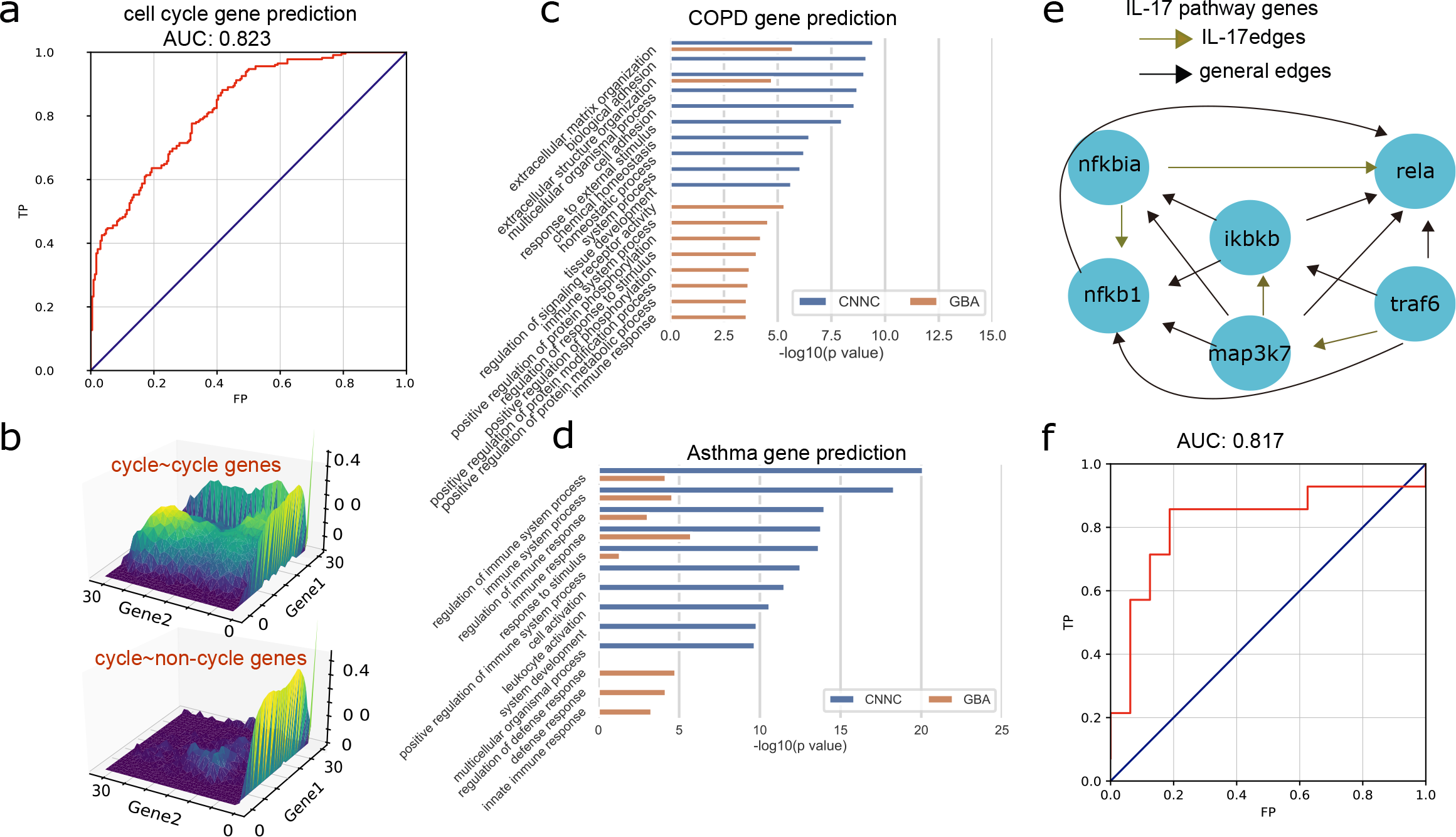
Functional assignment and pathway reconstruction using CNNC. CNNC can be used as a component in downstream analysis algorithms including for pathway analysis and functional assignments. (**a**) Performance of CNNC on the cell-cycle gene prediction task. (**b**) Top panel: predicted expression pattern of a cell-cycle~cell-cycle gene pair. Bottom panel: predicted expression pattern of a cell-cycle~non-cell-cycle gene pair. (**c, d**) The most significant GO terms of top 300 predicted COPD (**c**) and asthma (**d**) disease genes by CNNC and GBA respectively. (**e**) Directed edges annotated in KEGG for the IL-17 pathway gene nodes. (**f**) Performance for IL-17 pathway prediction task.

Given its success on a well-studied functional set we next asked if CNNC can be used to predict novel disease genes. We focused on two lung diseases, asthma and Chronic obstructive pulmonary disease (COPD). We obtained 147 and 44 genes for asthma and COPD respectively from ‘Malacards’^29^. We next trained CNNC with all known genes for each of the two diseases and used it to predict additional genes for each disease. We evaluated the predicted set both manually and by statistical analysis using GO and compared these to prior methods for Guilt By Association (GBA)^30^ analysis. As can be seem in **Fig. 5c** and **5d**, for both diseases, CNNC obtained much more significant GO terms when compared to GBA. Manual inspection of the top 10 genes for asthma indicated that 7 of them are supported based on recent studies (**Supplementary Tab. 3**) including ‘Lck’ which was recently determined to be a potential drug target for asthma therapy^31^.

### Applications of CNNC to pathway reconstruction

Given the results for KEGG we asked whether we can use CNNC to infer missing edges in current pathways. There have been several attempts to utilize expression and other data to further refine known pathways and many of these are based on co-expression analysis^6, 10, 32–34^. Since our method provides both direction and score we can extract all predicted directed edges above a certain score and compare the resulting pathway to the database pathway to see if any additional edges are predicted by our method. For this we focused on the interleukin 17 (IL-17) pathway from KEGG database, which plays crucial roles in inflammatory responses. We extracted 6 proteins and 4 directed edges from this pathway by only using directed edges with activation or inhibition edge types and filtering out cyclic gene pairs (**Fig. 5e**). The other 10 edges were not present in KEGG as causal interactions in IL-17 but were supported by other pathways. We applied CNNC trained on all database pathway edges that do not contain any of these 6 proteins. As can be seen (**Fig. 5f**), CNNC achieved a high AUROC of 0.82 for this task.

### Discussion and conclusion

Several methods for inferring gene-gene relationships from expression data have been developed over the last two decades. While these methods perform well in some cases, they suffer from a number of drawbacks that often led to overfitting (false positives) or missing key relationships (false negatives). The former can be attributed to the unsupervised nature of most methods (including methods for co-expression and clustering) making it hard to ‘train’ them on a labeled dataset. The latter often resulted from the assumptions used by specific methods (for example, distribution assumptions for DBNs) which do not always hold.

To address these issues we presented CNNC, a general framework for gene relationship inference which is based on convolutional NN (CNN). The key idea here is to convert the input data into a co-occurrence histogram which is very suitable for CNNs. Unlike most prior methods our method is supervised which allows the CNN to zoom in on subtle differences between positive and negative pairs. Supervision also helps fine tune the scoring function based on the different application. For example, different features may be important for analyzing TF-gene interactions when compared to inferring proteins in the same pathway. In addition to the supervised approach the fact that the network can utilize the large volumes of scRNA-Seq data allows it to better overcome masking issues reducing false negative.

Analysis of several different interaction prediction and functional assignment tasks indicates that CNNC can improve upon prior, unsupervised methods. It can also be naturally extended to integrate complementary data including epigenetic and sequence information. Comparisons to more advanced methods for biological network reconstruction further highlight the advantages of CNNC. In addition, CNNC can be used as a pre-processing step, or as a component in more advanced network reconstruction methods. Finally, CNNC is easy to use either with general data or with condition specific data (**Supplementary Fig. 10**). For the former, users can download the data and implementation from the supporting website, provide a list of labels (positive and negative pairs for their system of interest) and retrieve the scores for all possible gene pairs. These in turn can be used for any downstream application including clustering, network analysis, functional gene assignment etc.

CNNC is implemented in Python and both data and an open source version of the software are available from the supporting website.

## Online methods

### Dataset sources and pre-process pipelines

We used mouse scRNA-Seq dataset collected by Alavi et al^13^. The dataset consists of uniformly processed 43,261 expression profiles from over 500 different scRNA-Seq studies. For each profile, expression values are available for the same set of 20,463 genes. Among of the 43,261 cells, 2,696 are mESCs, 4,126 are dendritic cells, and 6,283 are bone marrow cells. mESC-time data which contains 3,456 cells was downloaded from GEO with accession number GSE79578^35^ and mESC-LIF data which contains 2,717 cells was downloaded from GEO with accession number GSE65525^36^. Mouse bulk RNA-Seq dataset were downloaded from Mouse Encode project^23^. That data included 249 samples and we only utilized genes that are present in the scRNA-Seq dataset leading to the same number of genes for both datasets. mESC Dnase data was also downloaded from Mouse Encode project^23^ (ENCFF096WRW.bed). Mouse TF motif information is from TRANSFAC database^37^. PWM values were calculated by Python package ‘Biopython’^38^.

For the DNase and PWM analysis we followed prior papers and defined the transcription start site (TSS) region as 10KB upstream to 1KB downstream from the TSS for each gene^19, 20^. For each TF and gene pair, using Biopython package we calculated the score between the TF motif sequence and both the ‘+/-’ sequences at all possible positions along the TSS region of the gene, and then selected the maximum one as the final PWM score. The maximum Dnase peak signal in the TSS region was calculated as the scalar Dnase value for each gene.

### Labeled data

mESC ChIP-seq peak region data was downloaded from GTRD database, and we used peaks with threshold p value < 10^−400^ for mESC cells and 10^−200^ for bone marrow cells and dendritic cells. If one TF X has at least one ChIP-seq peak signal in or partially in the TSS region of gene Y, as defined above, we say that X regulates Y. KEGG and Reactome pathway data were downloaded by the R package ‘graphite’^39^.

KEGG contains 290 pathways and Reactome contains 1581 pathways. For both, we only select directed edges with either activation or inhibition edge types and filter out cyclic gene pairs where genes regulate each other mutually (to allow for a unique label for each pair). In total, we have 3,057 proteins with outgoing directed edges in KEGG and the total number of directed edges is 33,127. For Reactome the corresponding numbers are 2,519 and 33,641.

### Constructing the input histogram

For any gene pair *a* and *b*, we first log transformed their expression, and then uniformly divided the expression range of each gene to 32 bins. Next we created the 32×32 histogram by assigning each sample to an entry in the matrix and counting the number of samples for each entry. Due to the very low expression levels and even more so to dropouts in scRNA data, the value in zero-zero position is always very large and often dominates the entire matrix. To overcome this, we added pseudocounts to all entries. We combined bulk and scRNA-Seq NEPDFs by concatenating them as a 32×64 matrix to achieve better performance.

### CNN for RPKM data

We followed VGGnet^40^ to build our convolutional neural networks (CNN) model (**Supplementary Fig. 1**). The CNN consists of stacked layers of *x* 3×3 convolutional filters (equation (1)) (*x* is a power of 2, ranging from 32 to 64 to 128) and interleaved layers of 2×2 maxpooling (equation (2)). We used the constructed input data as input to CNN. Each convolution layer computes the following function: 

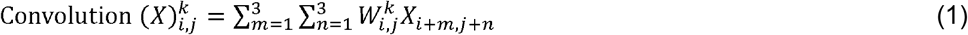

Where *X* is the input from the previous layer, (*i,j*) is output position, *k* is convolutional filter index and *W* is the filter matrix of size 3×3. In other words, each convolutional layer computes a weighted average of the prior layer values where the weights are determined based on training. The maxpooling layer computes the following function: 

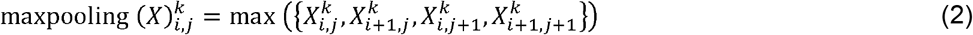

Where *X* is input, (*i,j*) is output position and *k* is the convolutional filter index. In other words, the layer selects one of the values of the previous layer to move forward.

### Overall structure of CNNC

The overall structure of the CNN is presented in **Supplementary Fig. 1**. The input layer of the CNN is either 32×32 or 32×64 as discussed above. In addition, the CNN contains 10 intermediate layers and a single one or three-dimension output layer. The ten layers include both convolutional and maxpooling layers, and the exact dimensions of each layer are shown in **Supplementary Fig. 1**. Following ref 41^41^ we used rectified linear activation function (ReLU) as the activation function (equation (3)) across the whole network, except the final classification layers where ‘sigmoid’ function (equation (4)) was used for two categories classification and ‘softmax’ function (equation (5)) for multiple categories classification. These functions are defined below. 

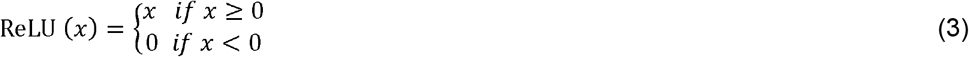

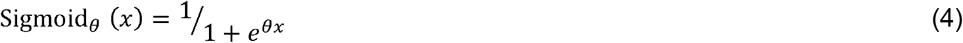

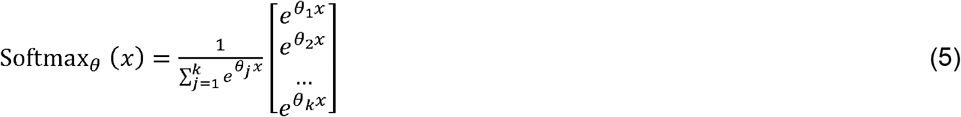

### Training and testing strategy

We evaluated the CNN using cross validation. In these, training and test datasets are strictly separated to avoid information leakage. See **Supplementary Methods**, and **Supplementary Table 1** for details. For the three labels (causality analysis) we did the following: for each gene, we generated (*a, b*) (label1) and (*b, a*)’s (label2) NEPDF matrices. For the 0 label we generated a (*a*, *N*) NEPDF matrices for GTRD where *N* was a random gene among all non-targets and *a* was the TF. 0 labels for KEGG or Reactome were generated from random (*M, N*) gene pairs among KEGG or Reactome gene sets. After training, we used *p*1(*a, b*) + *p*2(*a, b*) as the probability that *a* interacts *b*, *p*2(*a, b*) – *p*2(*b, a*) as the pseudo probability that *b* regulates *a*.

### Integrating expression, sequence and DNase data

To integrate Dnase and PWM data with the processed RNA-Seq data, we first computed the max value for a PWM scan and DNase accessibility for each promotor region. We next generated a two-value vector from this data for each pair and embedded it to a 512D vector using one fully connected layer containing 512 nodes. Next these are concatenated with the expression processed data to form a 1024D vector which serves as input to a fully connected 512-node plus 128-node layer neural network classifier. See **Supplementary Fig. 1** for details. Early stopping strategy by monitoring validation loss function is used to avoid overfitting.

### Functional gene assignment

To assign a function (biological process or disease involvement) we train a CNNC model for each known gene *g* for that function. Similar to all CNNCs, input to each model is a pair of genes where one is *g* and the other is either a positive (known) or negative gene. Next we built a two-layer fully connected neural network that take a vector of inputs (the 1 value output from each of the trained CNNC models) and outputs the final decision.

### Known genes for functional assignment testing

We downloaded 855 (182, 59) human cell cycle (asthma, COPD) genes from GSEA (‘Malacards’^29^ (a human disease website, https://www.malacards.org/)). We obtained mouse ontologies for all genes resulting in 682, 147 and 47 genes for cell cycle, asthma and COPD, respectively. For training we used all genes for the diseases and a randomly selected set of cell cycle genes. See **Supplementary Note** for details.

### Selection of edges for the IL-17 pathway analysis

We performed leave-one-pathway-out validation to evaluate CNNC’s performance for predicting edges for individual pathways. We selected a relatively small pathway (‘IL-17’ from KEGG) to improve our ability to present it visually. We discuss more general results for KEGG as well (**Fig. 4**). For this analysis we only selected directed edges with either activation or inhibition types and filtered out cyclic gene pairs where genes regulate each other mutually to purify the edge types. In total, we had 6 nodes and 4 directed edges for the IL-17 pathway. Next, we trained CNNC with the entire KEGG dataset excluding any interactions for the six ‘IL-17’ pathway proteins.

## Supporting information

SI table 3 for COPD

SI table 3 for asthma

## Data availability

All data, scripts and instruction required to run CNNC in Python can be found in our support website. All other public data can be found following the pipelines in “**Dataset sources and pre-process pipelines**” and “**Labeled data**” parts of Online methods.

## Acknowledgements

Work partially supported by NIH grant 1R01GM122096, US National Science Foundation (DBI-1356505) to ZBJ. and a James S. McDonnell Foundation Scholars Award in Studying Complex Systems to Z.B.-J.

## Author contributions

Y.Y. and Z.B.-J. conceived the method. Y.Y. implemented CNNC and the support website. Y.Y. and Z.B.-J. designed the experiments. Y.Y. and Z.B.-J. wrote the manuscript.

## Competing interests

None

