## Supplement for "Deep learning for inferring gene relationships from single-cell expression data"

### Supplementary results

#### Supplementary Figure 1

**CNNC’s architecture**

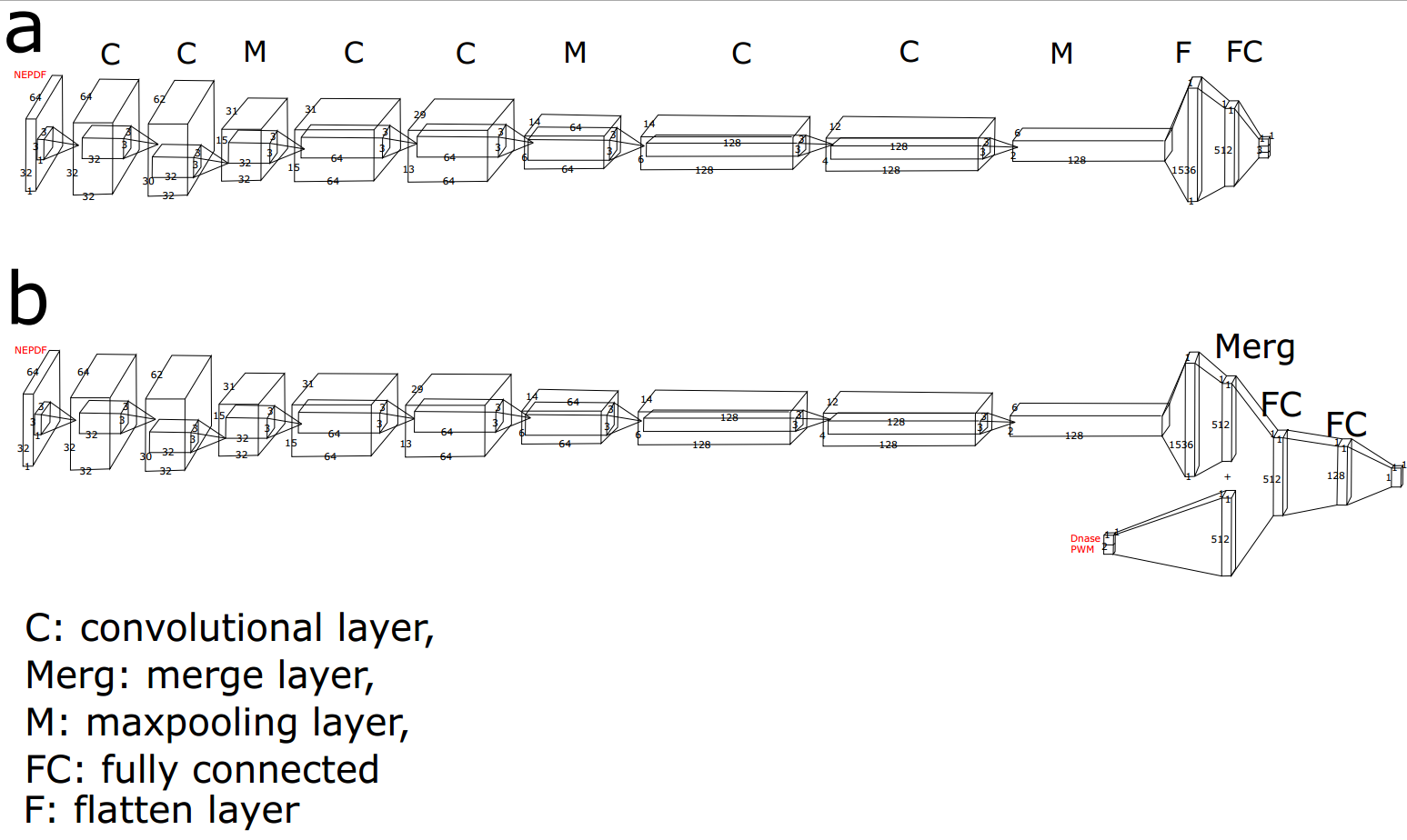

(**a**) Typical architecture of CNNC with combined NEPDF as input. The first layer is the input layer of 64×32×1, which is the exact size of NEPDF. Through 32 convolutional filters of 3×3×1 with padding technology, the input was transformed as a 64×32×32 tensor. Through 32 convolutional filters of 3×3×32 without padding technology, it was transformed as a 62X30X32 tensor. Through 2×2 maxpooling layer, the size of the tensor was transformed as 31×15×32. By such two stacked convolutional layers plus one maxpooling layer, the NEPDF was finally transformed as a 1536D vector through one flatten layer. Such vector was fed as a feature vector to a neural network classifier contained a 512-node hidden layer and 3-node output layer. (**b**) Structure of CNNC with NEPDF, PWM, Dnase-seq as input. Here the network embedding NEPDF as a 512D vector is identical to that in (**a**). The PWM and Dnase value was firstly organized as a 2D vector and then was embedded as 512D vector through a fully connected layer. The two 512D vectors were concatenated as a 1024D vector which serves as input to a neural network classifier with fully connected 512-node plus 128-node hidden layer plus 1-node output layer.

#### Supplementary Figure 2

**GTRD TF-target prediction based on five datasets by five methods**

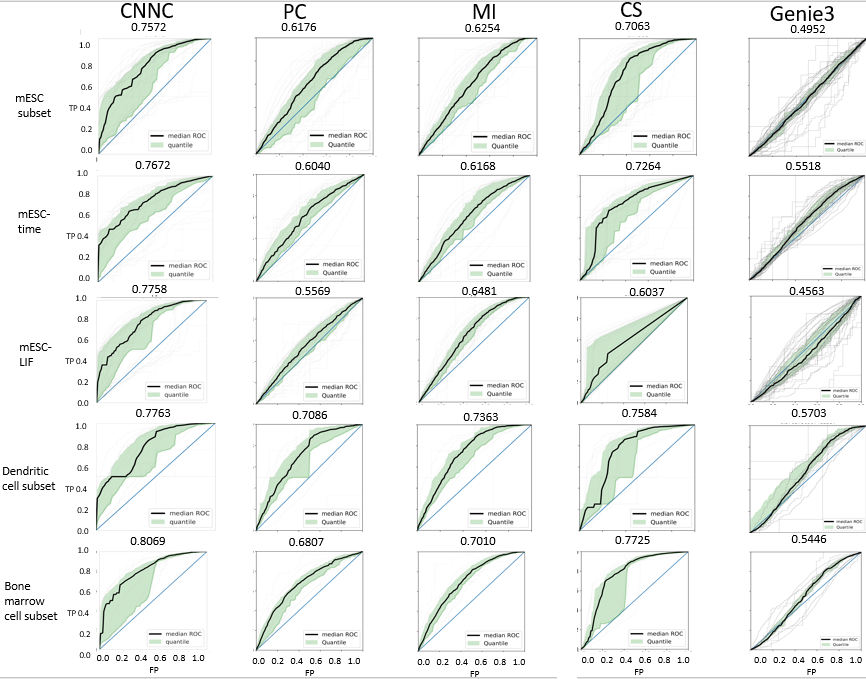

The column corresponds to CNNC, PC, MI, CS and Genie3 method respectively, each row corresponds to mESC subset, mESC-time, mESC-LIF, dendritic cell subset and bone marrow cell subset respectively. Light gray lines represent the performance for each TF. Black line represents the median ROC, and light green region represents the 25~75 quantile.

#### Supplementary Figure 3

**Three-fold cross validation for KEGG and Reactome database interaction (pathway edges) prediction by CNNC**

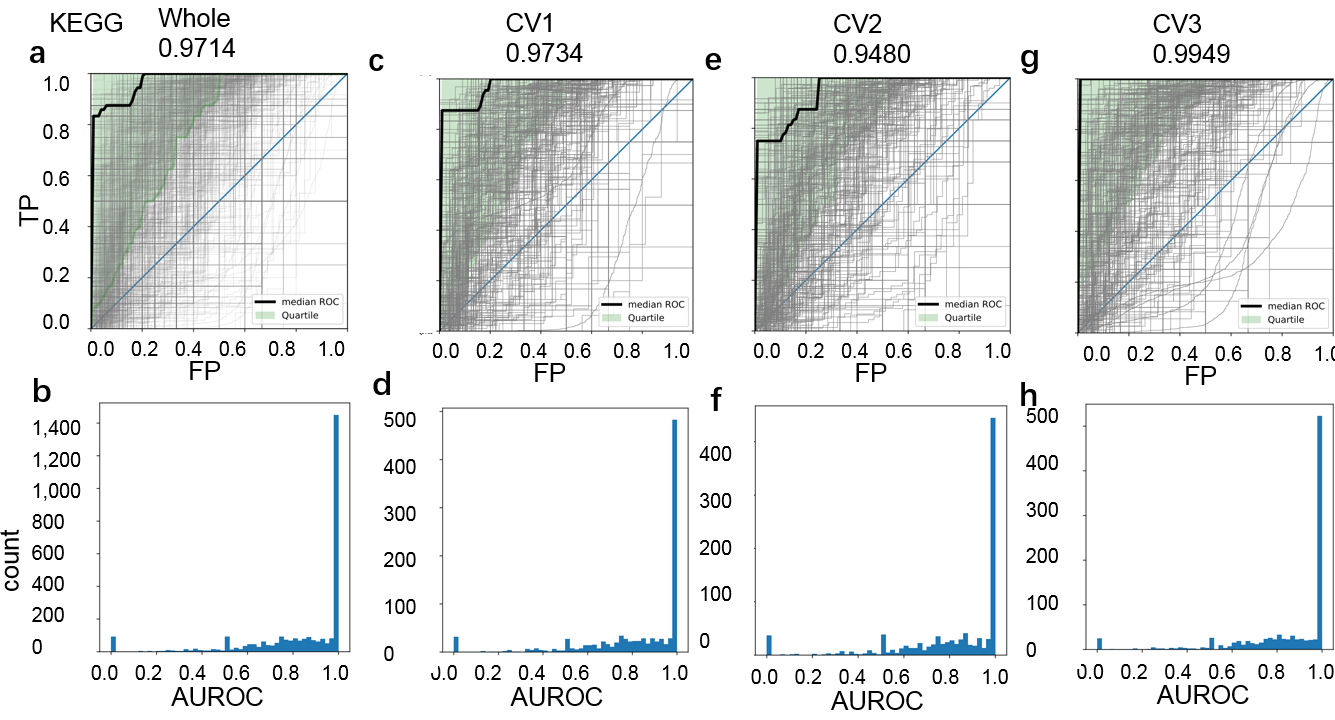

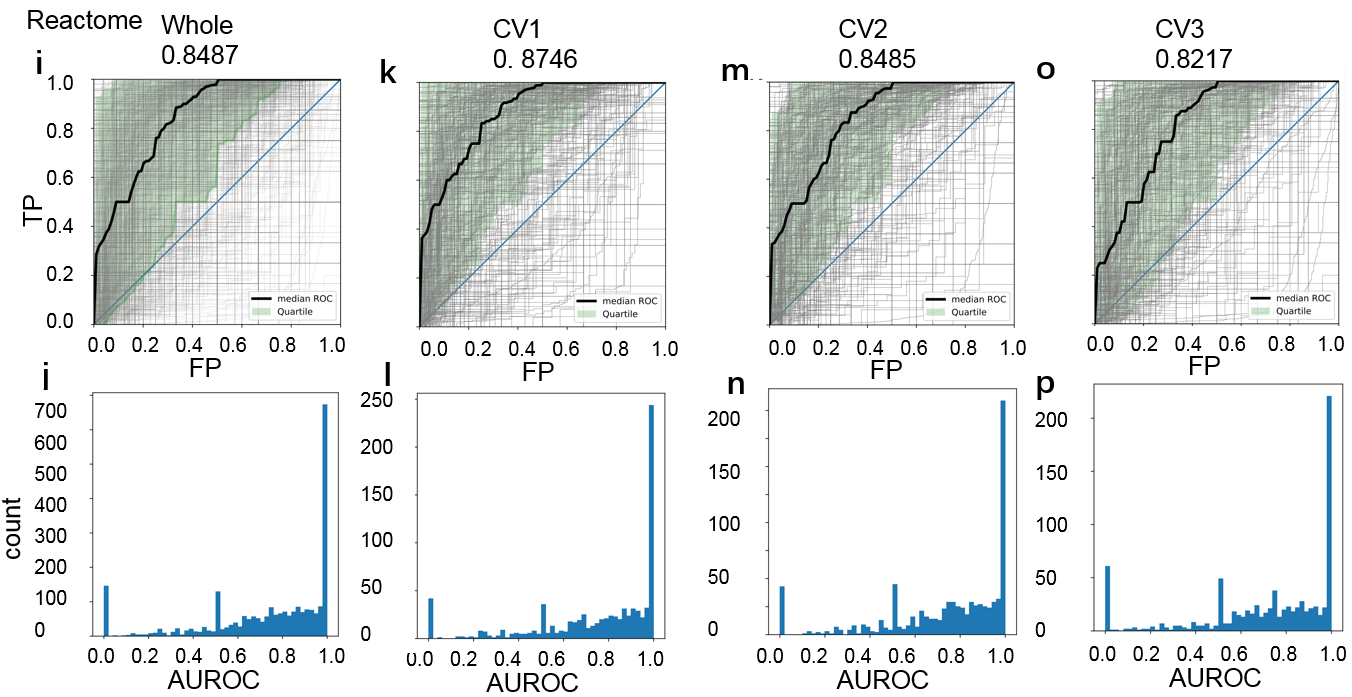

(Top) result for KEGG database. (Bottom) result for Reactome database. (**a**) The overall ROCs for the whole three-fold cross validation of KEGG. (**b**) The AUROC histogram of (**a**). (**c**) Fold 1’ ROCs. (**d**) The AUROC histogram of (**c**). (**e**) Fold 2’ ROCs. (**f**) The AUROC histogram of (**e**). (**g**) Fold 3’ ROCs. (**h**) The AUROC histogram of (**g**). (**i**) The overall ROCs for the whole three-fold cross validation of Reactome. (**j**) The AUROC histogram of (**i**). (**k**) Fold 1’ ROCs. (**l**) The AUROC histogram of (**k**). (**m**) Fold 2’ ROCs. (**n**) The AUROC histogram of (**m**). (**o**) Fold 3’ ROCs. (**p**) The AUROC histogram of (**o**).

#### Supplementary Figure 4

**Reactome database interaction prediction by CNNC**

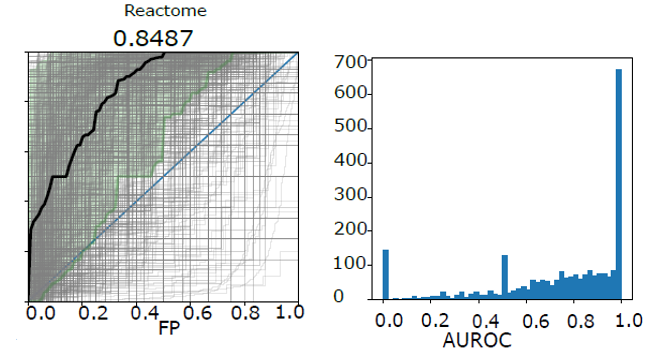

Left panel: Overall ROCs for CNNC performance on Reactome pathway gene interaction prediction with bulk and scRNA-Seq. Right panel: The Area Under the Receiver Operating Characteristic curve (AUROC) histogram for the left panel.

#### Supplementary Figure 5

**KEGG database interaction prediction by fully connected NN and CNN without pooling layer**

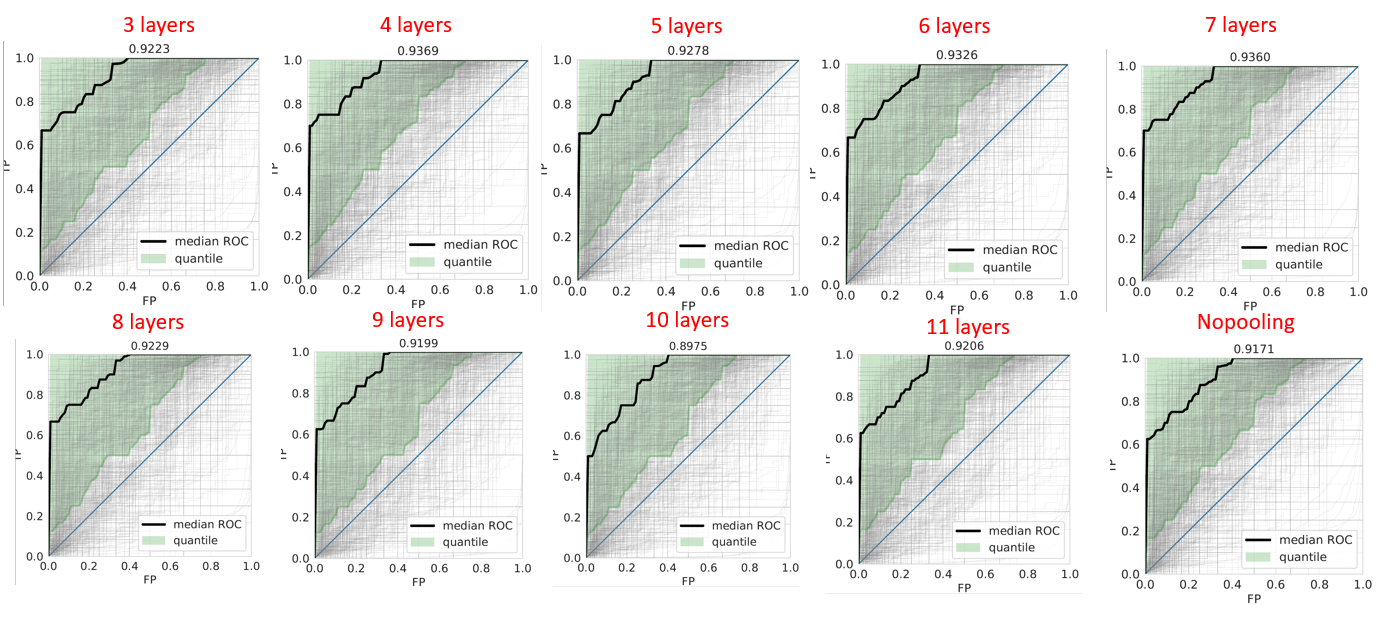

Here all neural networks’ input is 2048-dimension (32×64) vector reshaped from the NEPDF image. 3-11 layers of 512 nodes are used to build fully connected neural networks. CNN architecture with pooling layer was also implemented.

#### Supplementary Figure 6

**Causality prediction for GTRD, KEGG and Reactome database**

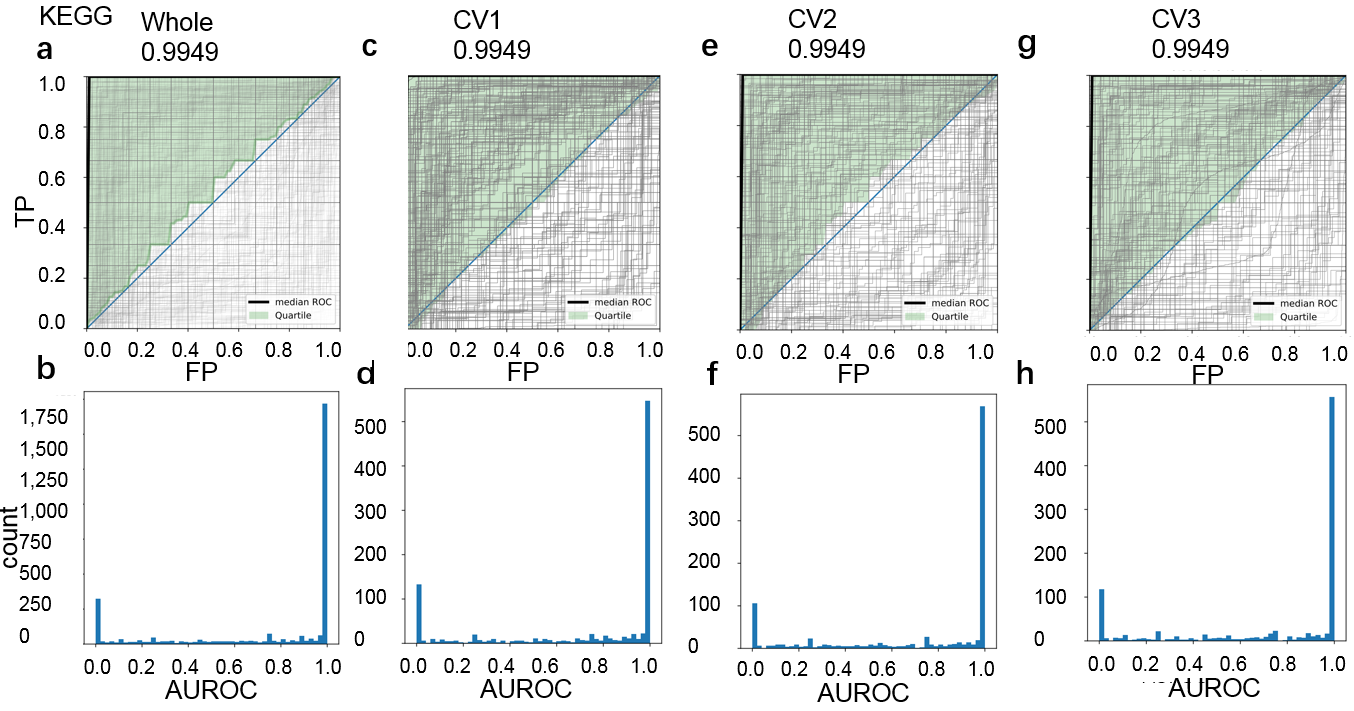

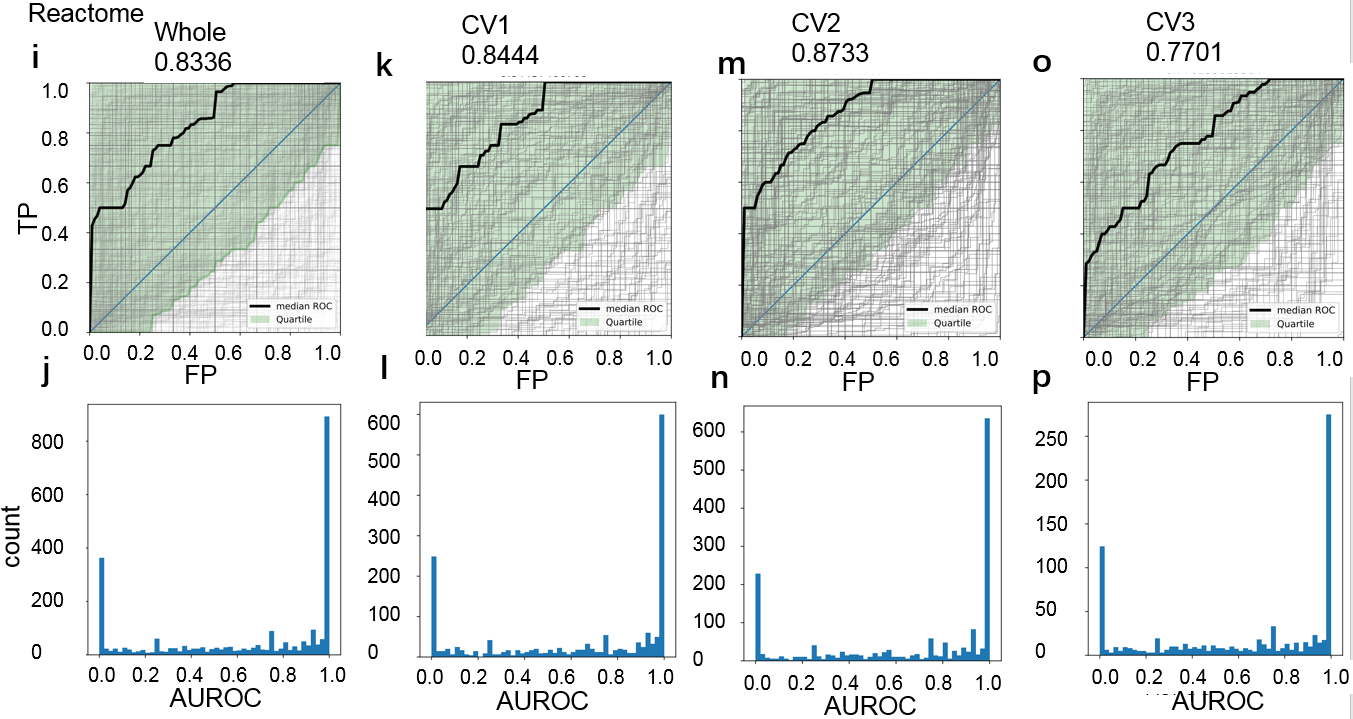

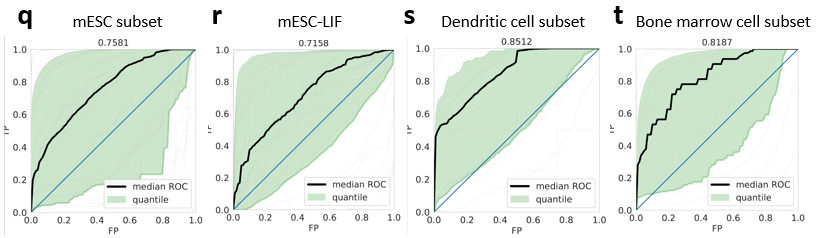

(**a-p**) **Three-fold cross validation for KEGG and Reactome database causality prediction by CNNC**. (Top) result for KEGG database. (Bottom) result for Reactome database. (**a**) The overall ROCs for the whole three-fold cross validation of KEGG. (**b**) The AUROC histogram of (**a**). (**c**) Fold 1’ ROCs. (**d**) The AUROC histogram of (**c**). (**e**) Fold 2’ ROCs. (**f**) The AUROC histogram of (**e**). (**g**) Fold 3’ ROCs. (**h**) The AUROC histogram of (**g**). (**i**) The overall ROCs for the whole three-fold cross validation of Reactome. (**j**) The AUROC histogram of (**i**). (**k**) Fold 1’ ROCs. (**l**) The AUROC histogram of (**k**). (**m**) Fold 2’ ROCs. (**n**) The AUROC histogram of (**m**). (**o**) Fold 3’ ROCs. (**p**) The AUROC histogram of (**o**). (**q-t**) causality prediction results for mESC subset, mESC-LIF, dendritic cell subset and bone marrow cell subset respectively.

#### Supplementary Figure 7

**CNNC VS down sampled BDN on KEGG dataset**

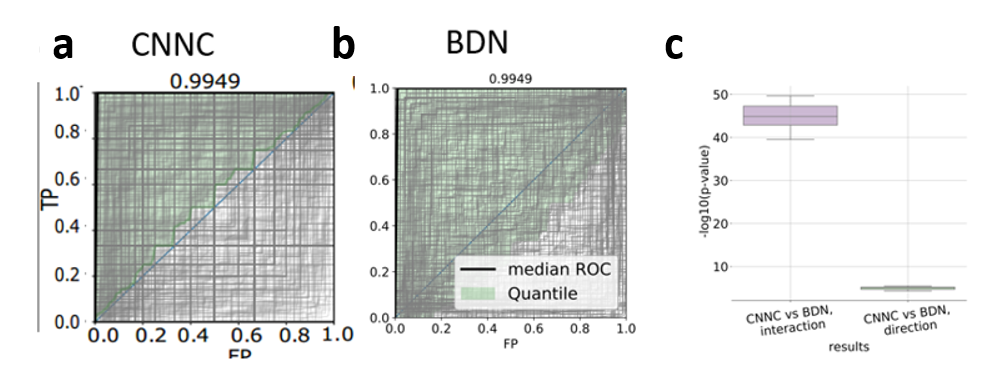

(**a, b**) CNNC and **down sampled** BDN’s performance on KEGG causality prediction. (**c**) Comparison between CNNC and downsampling BDN on KEGG interaction and causality prediction. 1000 samples were randomly selected from the whole data 10 times. P-values were calculated by pairwise wilcoxon test 10 times for each comparison (a total of 10 comparisons (10 p-values) for each column in **c**). Boxplot was shown with median, first, third quartile, maximum and minimum.

#### Supplementary Figure 8

**Down sampled CNNC VS down sampled BDN on KEGG dataset**

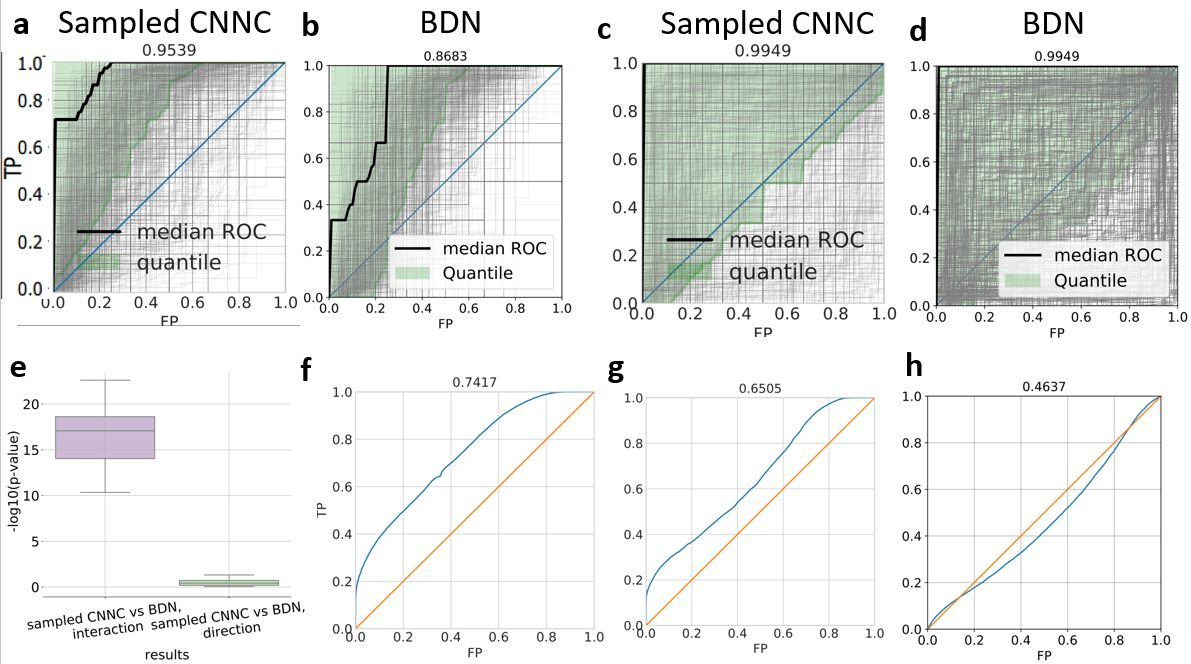

(**a, b**) CNNC and BDN’s performance based on downsamplings with size of 1000 on KEGG interaction prediction, which have been shown in Fig. 3. (**c, d**) CNNC (has been shown in Fig. 4) and BDN’s performance based on dowmsamplings with size of 1000 on KEGG causality prediction. (**e**) Comparison between CNNC and BDN on KEGG interaction and causality prediction. 1000 samples were randomly selected from the whole data 10 times. P-values were calculated by pairwise two-sided wilcoxon test 10 times for each comparison (totally 10 comparisons (10 p-values) for each column in **e**). (**f, g, h**) ROCs of the whole causality prediction without separation (in contrast to **Supplementary Fig.7a, Supplementary Fig. 8c and 8d** where the ROCs were separated by genes with outgoing edges).

#### Supplementary Figure 9

**KEGG database directed** **excitatory and inhibitory edges prediction by CNNC**

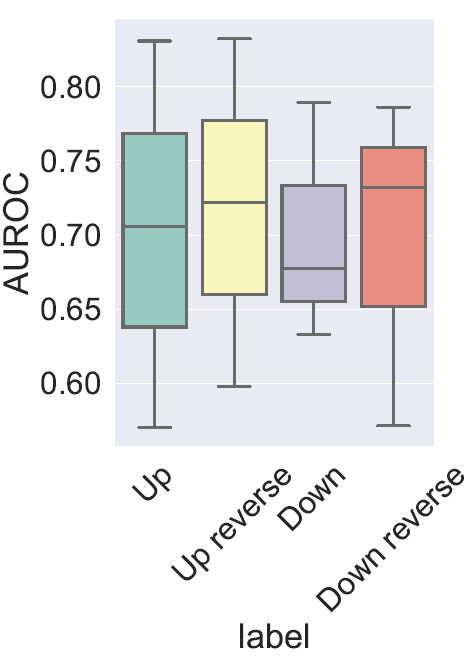

Here CNNC was trained by 4-category labels where label 1 means gene 1 activates gene 2 (Up), 2 means gene 2 is activated by gene 1 (Up reverse), label 3 means gene 1 represses gene 2 (Down) and label 4 means gene 2 is repressed by gene 1 (Down reverse).

#### Supplementary Figure 10

**Software pipelines for users**

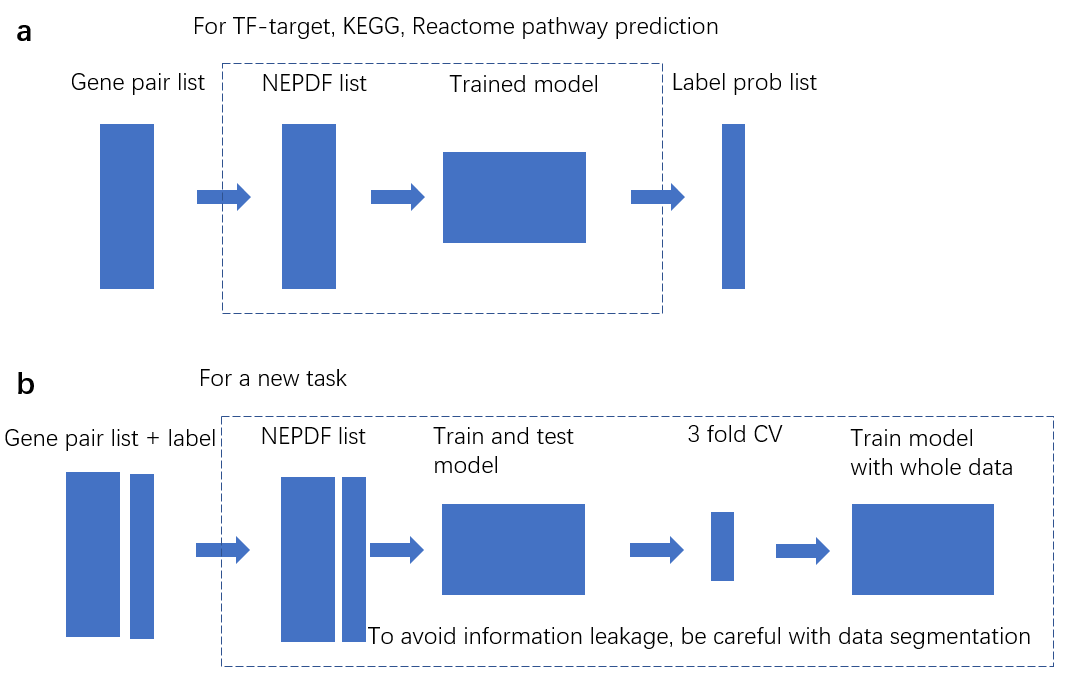

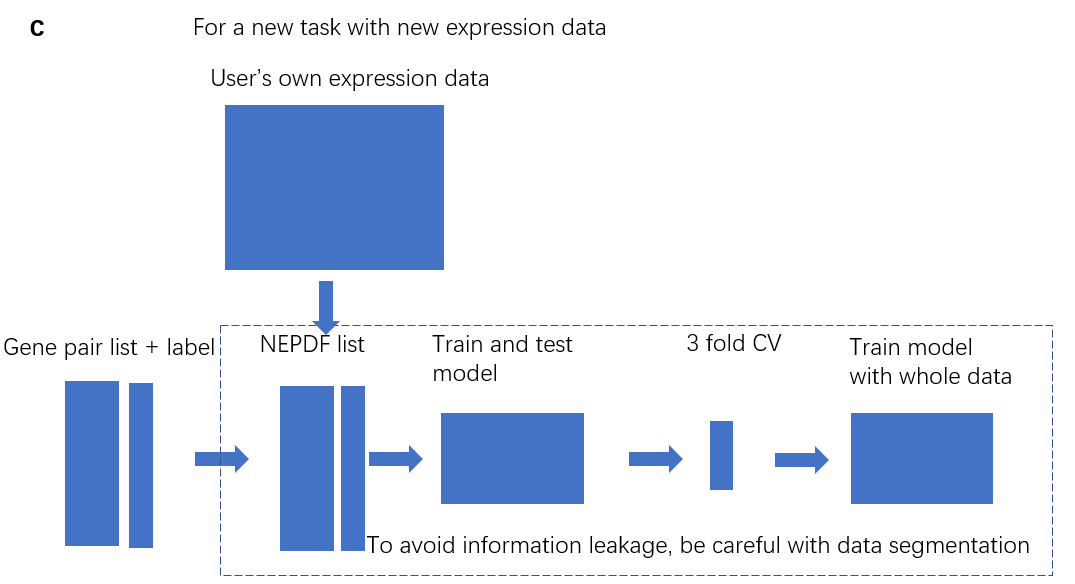

(**a**) Pipeline for TF-target, KEGG and Reactome edge predictions. Users only need to provide gene-pair candidate list. (**b**) Pipeline for a new task with the expression data we collected. Users need to provide gene-pair candidate list to generate NEPDF list and label list to train and test model. (**c**) Pipeline for a new task with the expression data users collect. Users need to provide gene-pair candidate list, their own expression data to generate NEPDF list, and label list to train and test model.

#### Supplementary Table 1

**Run time of each CNNC training process**

|  | Run time per epoch (*S*) | Max epochs | Patience |
| --- | --- | --- | --- |
| Fig. 2a, f, g, i, j, k | 20~100 (depends on size of training set) | 20 | None |
| Fig. 2p | ~20 | 200 | 100 |
| Fig. 3a | ~50 | 200 | 50 |
| Fig. 4 a/c/e | ~100 /~50 / ~60 | 20/200/200 | None/50/50 |
| Fig. 5a/c/e/f | ~1×454 networks/~2-4/~20-50/~83 | 200/200/200/100 | 80 |

In practice the actual number of epochs is always less than the max because of early stopping. Note that for **Fig. 2p** we only used a subset of the TFs used for **Fig. 2a** (19 of 38 TFs) for which we had motif information from TRANSFAC.

#### Supplementary Table 2

**Runtime of comparison methods**

| Methods | GTRD (cell-type specific data) | KEGG (the whole expression data) |
| --- | --- | --- |
| BDN  (interaction and direction) | Too large search space and too long time  × | ~ 1 hour for 1000 samples (8 cores, 16 GB). Terminated after 2,400 core hours without finishing for the whole 43,510 samples. |
| Count statistics  (interaction) | ~10 hours (1 core, 10 GB) | ~ 5 hours (1 core, 10 GB) |
| Genie3  (interaction) | ~4 hours (1 core, 15GB, for 38 TF models) | ~6 hours (1 core, 10 GB) |
| CNNC  (interaction and direction) | ~10 hours (38-fold CV (leave-one-TF-out), 1 core, 1 GPU, 20GB) | ~6 hours (three-fold CV, 1 core, 1 GPU, 20GB) |

×, did not finish in 2400 core hours; CV, cross validation.

### Supplementary Notes

#### Cross validation strategy

The general cross validation steps are as follow: The dataset is divided into three subsets, training set, validation set and test set. Training set is used to train the model with mini-batched stochastic gradient (MBSGD) algorithm, validation set is used to set hyperparameters and to determine the number of training iterations that are best for this data (to avoid overfitting) and test set is used for final performance evaluation.

In general, we fixed the total number of training iterations (Table. S1). We used a validation set to monitor the loss function (accuracy) and terminated training when the loss function (accuracy) began to increase (increase). See the supporting website for early stopping details. Run times are presented in **Supplementary Table 1**.

#### Comparison with Bayesian directed network (BDN), Count statistics and Genie3

##### BDN on KEGG interaction and direction prediction

Bayesian Decision Networks^1^ (BDN) were used in the past to infer pairwise interactions between genes. To compare the performance of CNNC with BDNs we constructed such networks from our data. For this, we relied on Maathuis et al.’s method^2, 3^ that generates the networks as follows: First it determines network skeletons by conditional independence test using PC algorithm. Next, it orients as many edges by V-structures as possible. Finally, it calculates casual effect from gene *a* to *b* as the final score for a given gene pair (*a, b*).

Through R package ‘pcalg’, BDN took more than 2,000 core hours to get the directed skeleton using the KEGG only genes (3,747 genes on Intel Xen W-2123 with 16GB memory). To shorten the runtime, we randomly downsampled 1000 samples from all 43,510 expression profiles 10 times, and then fed them to BDN PC algorithm. The skeletons contain around 10,000 edges, which is very sparse compared to the whole 3,747×3,747 possible edges.

The causal effect function ‘ida’ in R package ‘pcalg’ took more than two core hours for just one gene pair, which made BDN still impractical. To further shorten runtime, we implemented ‘ida’ algorithm using python without structure checking because the inferred skeleton was very sparse. Then the absolute value of casual effect was calculated as the final score for a given gene pair. When evaluating BDN’s performance, we selected the maximum between scores of (*a, b*) and (*b, a*) as the score of interaction prediction, and used the difference between scores of (*a, b*) and (*b, a*) as the score of direction prediction. We also trained CNNC based on the 1000 downsampled profiles (**Supplementary Fig. 8**).

##### Count statistics for KEGG and GTRD interaction prediction

Count statistics^4^ is a pair-wise method that directly calculates a score based on one gene pair’s expression profile. We used default parameter values setting K to 11, which was guided by the logarithm value of sample size to base ‘*e*’ and then used the ‘incSubseq_large_n.R’ function to calculate gene co-expression measure *W_2_* for our bulk and scRNA-seq data as non-time-series data.

##### Genie3 for KEGG and GTRD interaction prediction

Genie3^5^ uses a regression-like model to infer interactions. For each gene, Genie3 generates a tree-based ensemble model to predict its expression using other genes’ expression, and then ranks all the gene-pair coefficients. For the KEGG data, we first built 3,747 models each with 3,746 genes to predict the left one gene. However, such strategy took more than 200 core hours to calculate just one model. Since the original paper claimed that using normalized gene expression with unit variance, the weights from different models become comparable, we calculated the coefficients and ranked them for KEGG and GTRD data using the following strategies:

For KEGG data, each gene has a list of gene parents based on the gene pair list extracted from KEGG database. For each gene, we built a model predicting its expression using its parent genes’ expression. Since the coefficients from different model are comparable, we concatenated the whole gene-pair coefficients from all models together.

For GTRD data, each gene would have at more 38 TFs. For each gene, we built a model predicting its expression using all the 38 TFs and then concatenated all gene-pair coefficients together.

##### Cell-cycle gene, disease gene training and test data generation

We downloaded 855 (182, 59) human cell cycle (asthma, COPD) genes from GSEA^6^ (‘Malacards’^7^ (a human disease website, https://www.malacards.org/), ‘Malacards’) website, among which the overlap with mouse gene expression dataset is 682 (147,47) genes. We used 454 (98, 31) of them as ‘Training-known gene set’ for training and the left 228 (49,16) genes as ‘Test-known gene set’ for test. We randomly selected 454 (98,31) genes as ‘Training-unknown gene set’ and 228 (49,16) genes as ‘Test-unknown gene set’ from the whole gene set excluded the 682 (147,47) genes. To generate training gene pair set, we calculated NEPDFs for all gene pairs where gene 1 is in ‘Training-known gene set’ and gene 2 is in either ‘Training-known gene set’ (labeled as 1) or ‘‘Training-unknown gene set’’ (labeled as 0), thus got 454×454×2 (98×98×2, 31×31×2) gene pairs. To generate test gene pair set, we calculated NEPDFs for all gene pairs where gene 1 is in ‘Training-known gene set’ and gene 2 is in either ‘Test-known gene set’ (labeled as 1) or ‘‘Test-unknown gene set’’ (labeled as 0), thus got 454×228×2 (98×49×2, 31×16×2) gene pairs.

##### Cell-cycle gene training and test

During training, for each gene in ‘Training-known gene set’ which has 454×2 gene pairs, we trained a CNNC model, thus got 454 independent CNNC networks. Next we built a two-layer fully connected neural networks with 454-node input and 1-node output to ensemble all 454 CNNC networks. All training processes are based only on training set. Then the jointed NN model was evaluated on the test data.

##### Human disease gene training, test and comparison

First, we did a three-fold cross validation training to find the best training stop (training accuracy) where test accuracy was maximum. Then we generated a ‘whole known diseases gene training set’ that gene1 is in ‘Training-known gene set’ + ‘Test-known gene set’ (147, 47) and gene2 is in ‘Training-known gene set’ + ‘Test-known gene set’ +‘Training-unknown gene set’ + ‘Test-unknown gene set’ (147×2, 47×2), and we also generated a ‘whole unknown gene test set’ that gene1 is in ‘Training-known gene set’ + ‘Test-known gene set’ (147, 47) and gene2 are all genes never involved in the training (# of all left genes). During training CNNC with ‘whole known diseases gene training set’, we stopped the training when the training accuracy reached the best point we found before and then used CNNC to get scores for the ‘whole unknown gene test set’. As a result, each candidate gene has scores of a constant number (147, 47), we then sum them up as the final score by CNNC. GBA^8^ was implemented as follow: for each candidate unknown gene, we calculated the Pearson correlation between it and all known disease genes (147, 47) then sum them up as the final GBA score of this gene. Finally, we submitted the top 300 ranked predicted gene list to do GO term^9^ analysis and following comparison.
